# Human gut mobileome during antibiotic therapy. A trajectory-based approach to analysis of the metagenomic time-series datasets

**DOI:** 10.1101/793737

**Authors:** Anna Górska, Robert Schlaberg, Matthias Willmann, Jan Liese, Julia Victoria Monjaras Feria, Samuel Wagner, Ingo Autenrieth, Daniel H. Huson, Silke Peter

## Abstract

**Motivation:** Antibiotic resistance is widely recognized as a severe threat to current medical practice. Each antibiotic therapy drives the emergence and subsequent retention of antibiotics resistance genes within the human gut microbiome. However, the details on how the resistance spreads between bacteria within the human gut remain unknown, as does the role of horizontal gene transfer in this process, too.

**Results:** We present a novel approach to the analysis of time-series whole-genome metagenomic sequencing data. This involves partitioning the scaffolds from the metagenomic assembly into groups corresponding to bacterial chromosomes, plasmids and those with prophages and transposons. Using specialized sequencing of the bacteriophages we were able to track the flow of ciprofloxacin resistance genes from bacterial chromosomes, through the plasmids, to prophages and phages.

## 1 Introduction

Antibiotics are one of the most critical and successful classes of drugs in the history of medicine (Martinez, 2009). However, growing antibiotic resistance is recognized as a severe threat to modern medical practice by global agencies such as the World Health Organisation (WHO, 2014). It is established that the antibiotic usage drives emergence of antibiotic resistance (World Health Orginisation, 2017). Currently, metagenomic studies of the human gut microbiome provide insights into the microbiome’s response to antibiotic therapy and resistance emergence.

Therapy with antibiotics strongly disturbs the patient’s gut microbiome, in terms of both taxonomic and functional profiles (Huse *et al.*, 2008; Modi *et al.*, 2014). However, a *healthy* gut microbiome should be able to restore its taxonomic structure up to six months after the therapy (Raymond *et al.*, 2016). Yet, in a functional sense, each singular antibiotic therapy has a long-term impact on the patients’ gut microbiome.

Antibiotic treatment interacts with the gut bacteria in two ways. It selects the resistant bacteria and exerts environmental pressure promoting horizontal gene transfer (Broaders *et al.*, 2013). Those two phenomena lead to a permanent increase in the number and diversity of antibiotic-resistant genes. This, in turn, worsens the microbiome’s response to future antibiotic therapies (Penders *et al.*, 2013; Schaik, 2015; Francino, 2016).

The gut microbiome is a dynamic network of bacteria and bacteriophages, connected by occurrences of horizontal gene transfer (HGT). Using metagenomic sequencing, we can estimate the relative abundance of bacterial genomes and mobile genetic elements (MGEs) such as plasmids, phages, and transposons. Antibiotic treatment impacts the bacterial cells, and therefore the abundance of the chromosome, plasmid, and transposon sequences, but not directly those of the phages. Although resistance emergence in response to antibiotic therapy in the gut microbiome has been reliably observed, detailed mechanisms have not yet been described.

We studied the role of HGT in the emergence of antibiotic resistance within the gut microbiome of two healthy volunteers throughout a six-day ciprofloxacin therapy. To analyze the phages in the gut microbiome, phage-only sequencing (Phageome) was carried out alongside the standard whole-genome microbiome sequencing (Microbiome) for the stool samples.

In Górska *et al.* (2018) we described the analysis of these data focused on the Phageome. Here, we provide a more detailed analysis including machine learning approaches (Random Forest). We have divided the assembly of the Microbiome samples into bacterial chromosomes and MGE classes, and show the transfer of the resistant genes between them. The data comprises information on only two participants, therefore the presented results are not statistically robust. However, using this example, we present a novel approach to bioinformatic analysis of such a time-series metagenomic sequencing datasets.

## 2 Methods

Twelve stool samples collected from two participants at six time points were each sequenced twice, using two protocols, namely an ultra-deep microbiome sequencing (Microbiome set) and the sequencing of the DNA-phage fraction (Phageome set). The samples were taken before the start of ciprofloxacin therapy (0^*th*^ day), during (1^*st*^, 3^*rd*^ and 6^*th*^ days) and two samples after the therapy ended (+2^*nd*^ and +28^*th*^ days). Details on the sequencing protocol can be found in Górska *et al.* (2018) and Willmann *et al.* (2015). In brief, virus-like particles (VLPs) were extracted following the protocol described in (Thurber *et al.*, 2009) including two filtration and an ultracentrifugation step. Library preparation was performed using the Nextera XT DNA Library Preparation Kit (Illumina), followed by sequencing on the NextSeq 500 System (2×150 bp). For the microbiome dataset, DNA was extracted according to the human microbiome project protocol, followed by sequencing at GATC Biotech AG (Constance, Germany) using an Illumina MiSeq system (2×300 bp). In total, 24 datasets underwent bioinformatics analysis.

The standard pipeline for analysis of such metagenomic datasets relies on reads alignment against NCBI-nr database (Huson *et al.*, 2016). Here we aimed at associating antibiotic-resistance genes with mobile genetic elements. MGEs do not carry any specific genes or other genetic markers, that enable their identification within a metagenomic dataset. Therefore, the analysis relied on metagenomic assembly followed by thorough annotation of the scaffolds. Here we discuss further developments of the previous pipeline including concepts and methods that enabled gene-level analysis of the dynamics of MGEs within a human gut microbiome. In all instances sequencing reads were used, those were the *cleaned reads* sets, with removed contamination sequences (as described in Górska *et al.* (2018)). Python (v. 3.5.2) was used for pipeline programming along with the Matplotlib package (Hunter, 2007) that was used for plotting.

### 2.1 Diversity measurements

Alpha diversity is often employed for the first general description of time-series metagenomic datasets. We used it to compare the general diversity trends in the Microbiome and Phageome sets. Phages have high diversity, and their genomes are under-represented in the databases. Therefore, taxonomy-based diversity measurements are inadequate in the case of the Phageome set. Accordingly, the diversity was measured based on the clustered scaffolds.

First, all scaffolds longer than 200 bp for all samples within a variant, defined by a participant and sequencing run, were clustered (≥ 90% sequence similarity) using CD-HIT (Li and Godzik, 2006). Next, the longest scaffolds within a cluster were extracted. Subsequently, the reads were mapped onto the cluster representatives for of each sample separately. Finally, the cluster representatives were treated as the unit of diversity, and the portion of reads mapped onto them as an abundance. For these data, the Shannon diversity index (Shannon, 1948) was computed. The diversity values were arranged by the time point within a variant so that the diversity trajectories for the Phageome and Microbiome sets could be plotted.

### 2.2 Annotation

In the analysis, we treated each of the scaffolds as a multidimensional data vector. The dimensions correspond to the various features, depicted in Fig. 1.

**Fig. 1.**
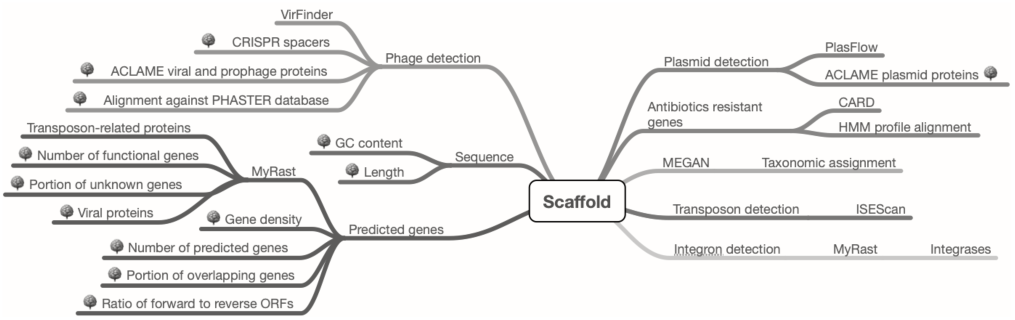
Features annotated for each of the scaffolds grouped by the category. Features marked with a tree icon were later used for phage prediction using a Random Forest.

First, the data vectors included essential characteristics, such as scaffold length, GC content, and abundance (short read coverage averaged by the scaffold length and the number of reads in the sample). Second, the vector contained features derived from the gene prediction (using prodigal (Hyatt *et al.*, 2010)), such as the number and density of genes and a portion of the reverse-oriented genes. The predicted genes were annotated based on the alignment to several databases, and separately by the functional annotation system MyRast (Overbeek *et al.*, 2014).

The third class of the features informed on the antibiotic resistance. Sequences of the predicted genes were aligned against two databases of antibiotic-resistant genes: CARD (Jia *et al.*, 2017) and ResFam (Gibson *et al.*, 2014). The first database was used for the protein-protein alignment, and only hits with high identity (≥ 80%) and coverage regarding length (≥ 80%) were accepted. The second database comprised HMM profiles of the resistant genes classes (alignment with HMMER (Eddy, 2011)). The HMM profile alignment provides higher sensitivity with lower specificity and allows alignments to the gene fragments.

Other features addressed the presence of MGEs, such as alignment to two specific databases of MGE proteins: ACLAME (Leplae *et al.*, 2009), and PHASTER (Arndt *et al.*, 2016), and scores provided by two k-mer based methods for predicting Plasmids and phages, PlasFlow (Krawczyk *et al.*, 2018) and VirFinder (Ren *et al.*, 2017) respectively. Finally, features included a taxonomic assignment by the MEGAN-LR pipeline (Huson *et al.*, 2018).

### 2.3 Identification of mobile genetic elements

Based on the annotations the Microbiome set was partitioned into bacterial chromosomes, plasmids, and scaffolds containing (pro)phages, or transposons. We assumed that the Phageme set contained only DNA of VLPs.

#### Phage identification

VirFinder (Ren *et al.*, 2017) was used for initial identification of phage scaffolds. VirFinder is trained based on k-mers extracted from the genomes of known phages, from the NCBI database (as of 2017). Therefore, its model does not encapsulate the entire space of the phage genetic diversity.

To enrich the phage scaffolds we trained a Random Forest (RF) classifier (Tin Kam Ho, 1995) using the Python sklearn package (Pedregosa and Varoquaux, 2011). The positive class consisted of all those scaffolds that had a low *p*-value in the VirFinder prediction. The data matrix comprised thirteen features representing the sequence, phage, and gene-related, and annotation-based features (labeled with a tree icon in Fig. 1).

Random Forest parameters were selected automatically using an inbuilt mechanism (final sklearn parameters: n_trees: 10000, max_depth: 5, max_features: 3, min_samples_split: 2). Overfitting was controlled with two tests. The first test measured advantage of the accuracy of the classifier’s prediction for the train set over the test set (*train* − *test*) across all runs. The second test measured portions of scaffolds denoted as phage by the entire RF set across a range of the cutoff values. The rationale is that better RF classifiers produce trajectories that plateau.

We used the Out-Of-Bag accuracy (OOB-accuracy) to evaluate the performance of the classifier, and mean decrease in accuracy to investigate feature importance. The classifier was trained on the under-sampled dataset containing the *positive* (phage) scaffolds and an equal number of the randomly chosen *negative* scaffolds (non-phage). Therefore, in each run, there is a large number of scaffolds not used for the training. Those are input in the newly trained classifier to predict whether a scaffold is a phage. The cycle was repeated 500 times so that for each *negative* scaffold the prediction was performed hundreds of times. Finally, we iterated through all *negative* scaffolds, and if the majority of the RF predictions were positive, we denoted a scaffold as a phage.

#### Plasmid identification

Plasmid scaffolds from the Microbiome set were defined as scaffolds, that did not contain phages (as defined above), were predicted as plasmid by PlasFlow (Krawczyk *et al.*, 2018) with a 95% cutoff and contained at least one protein annotated as plasmid in the ACLAME database (Leplae *et al.*, 2009).

#### Bacterial chromosome identification

Non-plasmid scaffolds, namely those that did not fulfill both criteria for plasmid, were denoted as bacterial chromosomes. Hence, the bacterial chromosome scaffolds can contain prophages and transposons.

#### Transposon identification

ISEScan (Xie and Tang, 2017) was used to annotate insertion sequences (IS). The scaffold was annotated as containing transposon if there were at least two ISes detected.

### Dynamics analysis

#### Scaffolds filtering

In this analysis, a single scaffold is represented by a multi-dimensional vector of features of different types. We can query features using various filters depending on the type of feature. The more complex features, such as taxonomic assignment, are hierarchically organized. The filter can be parametrized by the name of the feature, its level, and the value. The features can also be combined.

Such definition provides great freedom in choosing the groups of scaffolds. E.g., we could analyze a fraction of scaffolds denoted as a plasmid, carrying at least one resistant gene, and belonging to *Bacteroides* genus.

#### Feature-abundance trajectories

*Feature-abundance* was defined as a sum of the average coverage values for scaffolds that passed a defined filter. A *feature-abundance trajectory* is a vector of *feature abundance* at all consecutive timepoints. In the case of filters with hierarchical features, a decreasing number of scaffolds is used, and the higher level trajectories incorporate lower level trajectories.

We applied several trajectory levels to the analysis of this dataset. Starting from the most general using the most significant number of scaffolds, we progressed to the most precise filters, looking at the presence of the single genes.

The first level included a vast majority of the scaffolds and described a division between the main taxa affected by ciprofloxacin. Next, we analyzed trajectories for MGE selections and carriage of antibiotic-resistant genes. We also analyzed the trajectories for ciprofloxacin-resistant genes within the functional annotations.

#### Feature-abundance trajectory profiles

*Feature-abundance trajectory profiles* enable simultaneous analysis of the trajectories from the selected level of taxonomical and functional annotation methods. For taxonomic features, we analyzed genera, for functional assignments, we focused at genes. The average feature-abundance trajectories were extracted for all of the most detailed features. Three annotation methods were used for this step: taxonomic assignment (MEGAN), functional features (myRAST), and antibiotic-resistant genes (CARD).

The trajectories were normalized and clustered (using the Python scipy package AgglomerativeClustering with four clusters, complete linkage, cosine affinity). For each cluster, the *average scaled trajectory* was computed. Subsequently, the features were grouped based on the trajectory cluster. Next, we computed a *feature-abundance trajectory profile* reflecting portions of genes characterized by the most common trajectories.

The feature-abundance trajectory profiles differed depending on the underlying selection of the scaffolds. We analyzed the trajectory abundance for features amongst all scaffolds in the Phageome set and several scaffold groups in the Microbiome set, namely: bacterial chromosomes, plasmids, (pro)phage and transposon-carrying scaffolds.

## 3 Results

The number of input reads and portion of reads used in assembly are presented in Fig. S1. Sequencing and assembly statistics have been described before in (Górska *et al.*, 2018). Here we present the results of the extended analysis.

### Diversity trajectories

As expected, the Microbiome diversity decreased in response to the antibiotics therapy, from the 3^*rd*^ until the 6^*th*^ day, for both participants. It restored on the last day of sampling. The Phageome diversity displayed the opposite pattern. The diversity increased between the 3^*rd*^ and 6^*th*^ days of therapy and decreased on the last day in the case of participant A, and increased in the case of participant B (Fig. 2).

**Fig. 2.**
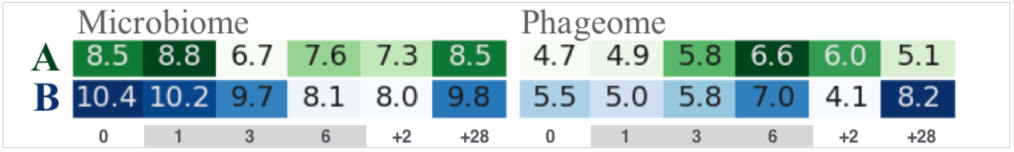
Diversity trajectories. The numbers and colors correspond to the Shannon index values. Colors are scaled separately for each trajectory. The gray bar indicates days of ciprofloxacin therapy.

The diversity measurements suggest that the changes are a response to the therapy. However, the response was shifted with respect to the therapy. Namely, the disturbance started on the 3^*rd*^ day of therapy and ended two days after the end of therapy (+2^*nd*^ day).

### Phage identification

Average OOB-accuracy of the RF runs with the best parameters was 69% for participant A and 68% for participant B, in both sets. In all runs, the number of MyRast functional genes, CRISPR spacers, and GC content were among the most influential features (Table S1). Feature selection, i.e., removing the features with low mean decrease in accuracy from the dataset, resulted in decreasing the accuracy. No single feature drove the scaffolds to be classified as phage. Hence the full set was used. In the Phage set, the CRISPR number and viral genes are differentiating. Whereas, in the Microbiome set, the viral genes of the ACLAME family and prodigal coverage were highly ranked.

### Mobile genetic element components

As predicted, the assembly of the Microbiome set consisted foremost of bacterial (∼ 90%), then (pro)phage, including both integrated phages and phage particles (∼ 10%), plasmid (∼ 2%), transposon (∼ 0.5%) and integron (∼ 0.3%) scaffolds. The break-down was similar between the participants. We assumed all scaffolds in the Phageome set were phages.

We assigned a known functional gene to ∼ 60% of scaffolds in both groups: the Phageome scaffolds and bacterial scaffolds in Microbiome. We detected a nearly complete antibiotic-resistant gene (CARD) on ∼ 0.3% of the scaffolds in Phageome and on an even smaller portion of the bacterial scaffolds. However, ∼ 5% of scaffolds in the Phageome and Microbiome sets had an alignment to a ResFam HMM profile, i.e., they carried a partial, or potentially unknown antibiotic-resistant gene.

### Feature-abundance trajectories

#### General taxonomic analysis

The taxonomic changes in the human gut microbiome prompted by the therapy with ciprofloxacin were characterized before (Stewardson *et al.*, 2015; Pérez-Cobas *et al.*, 2012). The authors reported that the abundances of Bifidobacterium, Faecalibacterium, Alistipes, Ruminococcus, and Dialister decreases in response to ciprofloxacin, whereas the abundance of Lachnospiraceae increases. This pattern was also observed for the Microbiome sets of the two participants (Fig. 3(a)).

**Fig. 3.**
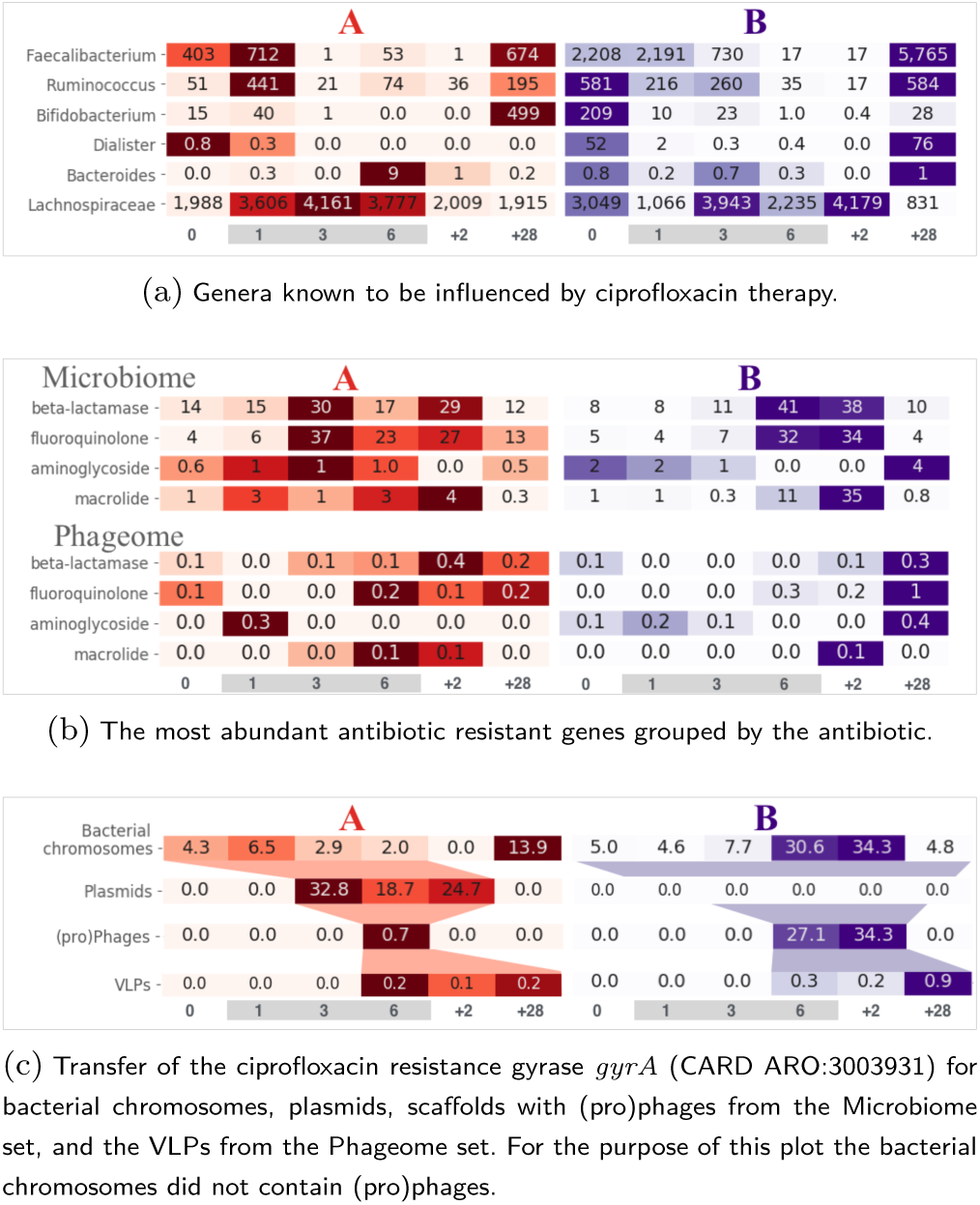
Feature abundance trajectories. The numbers and colors correspond to the feature abundance values. Colors suggest the general pattern of the trajectory, are assigned to one row at the time. The gray bars denote days of antibiotic therapy.

#### Antibiotic resistance dynamics

Among the antibiotic-grouped trajectories of antibiotic-resistant genes in the Microbiome set (Fig. 3(b)), fluoroquinolone and beta-lactams were the most abundant. In both participants, the resistance had *antibiotic-related trajectories*, i.e., their abundance increased between 3^*rd*^ and +2^*nd*^ days.

Virus-like-particles (VLPs) constituting the Phageome set also carried antibiotic-resistance genes. However, the increase of the trajectories of the two resistance classes started later in respect to the corresponding trajectories in the Microbiome set. That suggests that the phages picked up resistance after it was enriched within the bacterial cells. Moreover, the resistance genes persisted within the Phageome until the end of sampling.

#### Cirpofloxacin resistance

Resistance to ciprofloxacin can be conferred by a point mutation in the gyrase DNA, that inhibits binding by target alteration. Fig. 3(c) presents *feature abundance trajectories* for the most abundant gyrase conferring resistance to ciprofloxacin. The gene *gyrA* conferring resistance to fluoroquinolones was present on bacterial chromosomes in both participants. However, prompted by therapy with the antibiotic, it transferred onto plasmids, (pro)phages and phages in participant A. In the case of participant B, the resistance could not be detected in plasmids but in phages, indicating that a direct transfer from the bacterial chromosome might have occurred. Protein alignment and read mapping confirmed the mutation conferring resistance to the reference sequence (Fig. S2).

### Feature-abundance trajectory profiles

We identified ten patterns of trajectories, indicated by different colors in Fig. 4(a). First, there are trajectories that are flat or slightly fluctuating. Those correspond to genes, profiles or viral assignments with constant abundance. Next, there are trajectories with the strong dominance of a single day across the sampling. Such trajectories with the dominant maximum were divided into those that increase during the therapy-related days, so from 3^*rd*^ to +2^*nd*^ days, or outside them on the 0^*th*^, 1^*st*^and +28^*th*^ days.

**Fig. 4.**
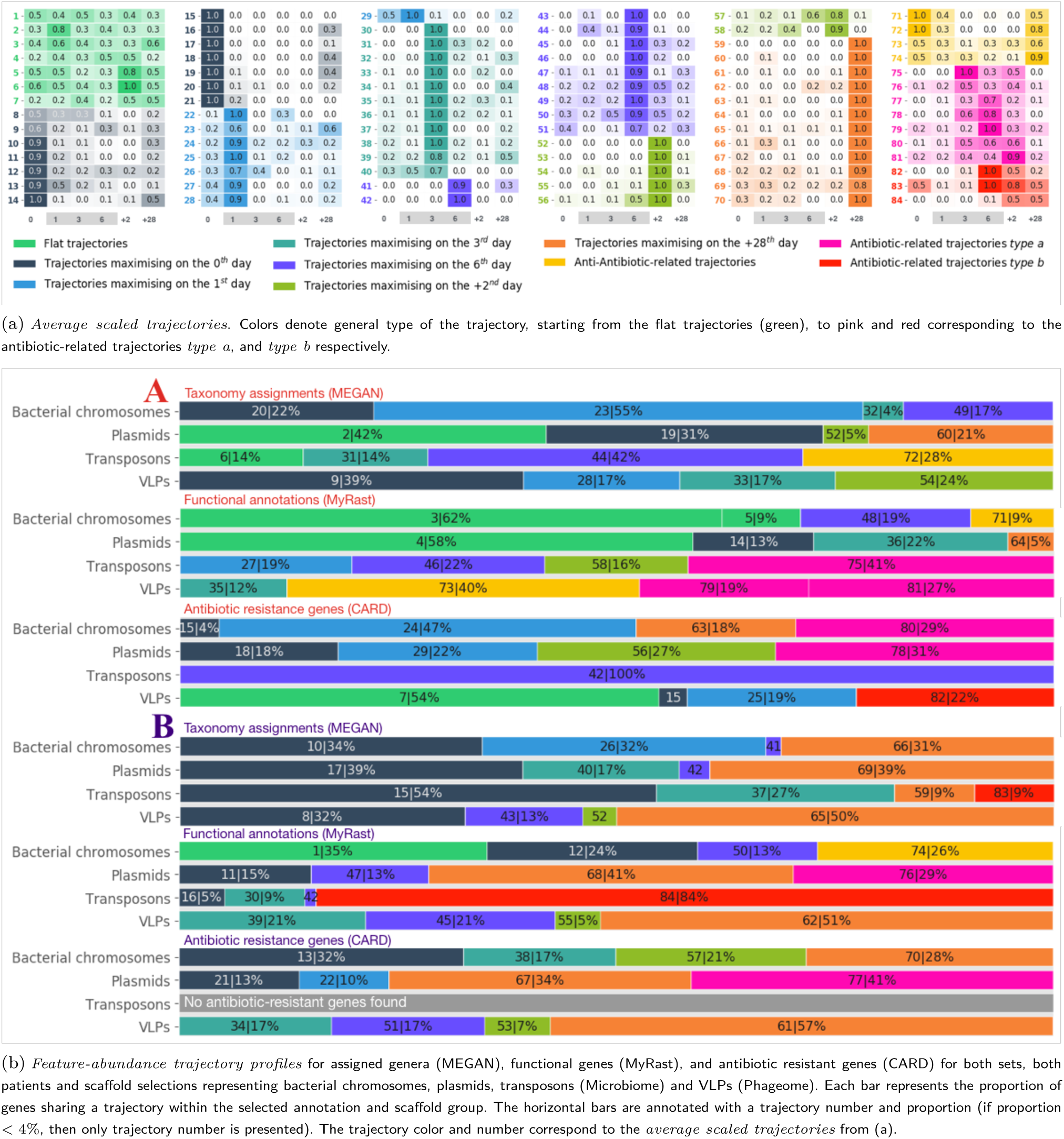
Global dynamics analysis.

We also identified three antibiotic-related trajectories. The first was characterized by an increase between the 3^*rd*^ and +2^*nd*^ days, analogous to the pattern revealed by the diversity trajectories (*type a*). The second antibiotic-related trajectory increased on the 6^*th*^ day of therapy and remained elevated until the end of sampling (*type b*). The last antibiotic-related trajectory decreased between the 3^*rd*^ and +2^*nd*^ days (Fig. 2).

All genes from CARD and myRAST as well as taxa assigned by MEGAN were annotated with a trajectory number. The table is available as a supplementary file (gene2trajectory.xlsx). The diversity measurements discussed above suggested that the participant B’s microbiome did not restore its structure after the antibiotic therapy. This is confirmed by the *feature-abundance trajectory profiles* analysis, as B’s profiles were dominated by the trajectories that maximized on the last day of sampling (Fig. 4(b)).

In this analysis, no genus annotated for bacterial chromosomes or plasmids displayed strictly antibiotic-related trajectories. Rather, a considerable portion (55% for A and 32% for B) of the bacterial-chromosomes genera had trajectories that were high in the first two time points and subsequently decreased but did not manage to restore their abundance within the 28 days of recovery.

The overall patterns of genus annotations differed from the patterns of the functional annotations for the same scaffold selections. The majority of functional genes (MyRast) on bacterial chromosomes (62% for A and 35% for B) had flat trajectories. That suggests that a change of the taxonomic structure does not necessarily cause an equivalent shift in the functional landscape.

The profile analysis co-discovered the pattern of the *gyrA* transfer. In participant A, the *gyrA* gene had a different trajectory for each of the scaffold selections corresponding to the MGEs (see supplementary gene2trajectory.xlsx). For plasmids, it had an antibiotic-related trajectory *type a* (number 78), for VLPs *type b* (number 82), and for the bacterial chromosomes a trajectory starting with low abundance and increasing on the last day of sampling (number 60).

A higher portion of antibiotic-related genes had antibiotic-related trajectories than of the overall functional genes. This portion was also more substantial for the MGEs. However, one has to keep in mind that the number of any annotated genes on the MGEs is lower than those of the bacterial chromosomes.

## 4 Discussion

The human gut microbiome can be analyzed from different perspectives using taxonomic or various functional classifications (Huson *et al.*, 2016). With the presented approach utilizing *feature abundance trajectory profiles*, all possible diverse features can be extracted and analyzed at once, providing a complex, but much more comprehensive view. This approach could be extended to a larger datasets of metagenomic samples. In that case, the number of trajectories and possibly their types would increase, but extracting groups of patients with similar *trajectory profiles* would be straightforward.

The *trajectory profiles* are only as informative as the underlying annotation methods are reliable. However, several annotation methods can be applied at once, and their results can be compared. Contrary to the classic approach, where the annotation methods have to be selected, here we can easily merge various sources, and extend the list of filters to create new selections.

Detection and analysis of mobile genetic elements (MGEs) within the human fecal metagenome sequencing data is a complex task, and the proposed MGE detection methods can surely be improved. Identification of MGEs requires information on several features of the sequence in the same time. Hence, machine learning methods are suited for solving this problem. We considered training the random forest classifier for identification of phage-carrying scaffolds using the Phageome set as ground truth, rather than the VirFinder results. The Phageome set was with 2×150 bp reads, compared to the Microbiome dataset for which 2×300 bp reads were obtained.

In this study, we demonstrated that the approach taken even enabled us to follow the dynamics of ciprofloxacin resistance on the level of point mutation in the *gyrA* gene. However, the entire analysis relies on the association of features, which is only possible for long scaffolds. Metagenome assembly can be error prone, due to the short sequencing reads and the unsaturated sequencing depth of the complex fecal samples.

Therefore the application of long read sequencing technology (e.g. Oxford Nanopore) in future studies, might enable more detailed plasmid analysis and phageome analysis without the need for independent sequencing and purification of VLPs (Bertrand *et al.*, 2018). However, the error rate so far is too high to use the k-mer based plasmid and phage detection methods.

## 5 Conclusion

The analysis of this small dataset shows that the presented approach enables both very detailed, and also global analysis of dynamics within the gut microbiome. To the best of our knowledge the approach is new. It was validated by a taxonomic analysis that aligned well with previous studies. We provide a *gene-level analysis* of the changes on the MGEs in the context of the entire human microbiome in a time-dependent manner in response to antibiotic the antibiotic therapy. We were able to show that specific antibiotic resistance genes transfer from the bacterial chromosomes, to plasmids and phages, where they persisted until the last day of sampling.

## Acknowledgements

The authors acknowledge support by the High Performance and Cloud Computing Group at the Zentrum für Datenverarbeitung of the University of Tübingen, the state of Baden-Württemberg through bwHPC and the German Research Foundation (DFG) through grant no INST 37/935-1 FUGG and grant no HU 566/12-1. We acknowledge support by the Open Access Publishing Fund of University of Tübingen.

