## Supplemental Figures and Tables for "Human gut mobileome during antibiotic therapy. A trajectory-based approach to analysis of the metagenomic time-series datasets"

### Supplementary Material

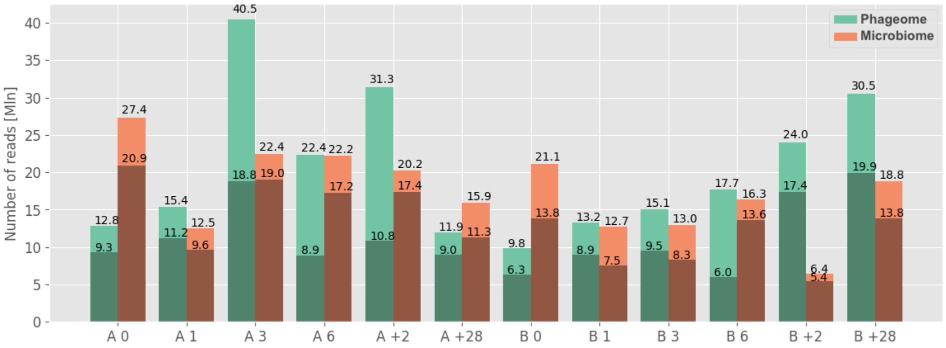


Figure S1: Number of reads per sample for both sequencing runs and particpants. The darker colors denote numbers of reads used for assembly.

Table S1: Mean decrease in accuracy for RF. Red and orange denote the highest and second highest values respectively within the variant.

| | Phageome | | Microbiome | |
|---|---|---|---|---|
| Feature | A | B | A | B |
| Number of MyRast functional genes | 26.5 ±2 | 26.2 ±2 | 26.9 ±1 | 15.9 ±1 |
| Scaffolds' GC-content | 15.0 ±2 | 23.3 ±2 | 16.9 ±2 | 29.6 ±1 |
| Scaffolds' length | 11.9 ±1 | 14.3 ±1 | 15.8 ± 1 | 20.7 ±1 |
| Number of CRISPR spacers | 13.4 ±2 | 12.6 ±1 | 10.5 ±1 | 4.5 ±1 |
| Number of viral genes (MyRast) | 10.0 ±2 | 6.0 ±1 | 2.4 ±1 | 5.3 ±1 |
| Portion of genes with unknown function | 5.0 ±1 | 6.2 ±1 | 4.7 ±1 | 2.2 |
| Number of viral genes (Phaster) | 6.4 ±1 | 4.4 ±1 | 0.0 | 0.0 |
| ACLAME classification (vir) | 4.3 ±1 | 1.0 | 0.0 | 0.0 |
| ACLAME classification (proph) | 1.8 ±1 | 0.0 | 0.0 | 0.0 |
| ACLAME classification (plasmid) | 0.9 | 0.4 | 0.0 | 0.0 |
| Predicted gene coverage | 0.0 | 0.0 | 6.3 ±1 | 5.0 ±1 |
| Number of predicted genes | 0.0 | 0.0 | 5.5 ±1 | 5.9 ±1 |
| Portion of reverse oriented ORFs | 0.0 | 0.0 | 1.9 ±1 | 2.9 ±1 |
| Portion of the overlapping genes | 0.0 | 0.0 | 2.2 | 1.05 |

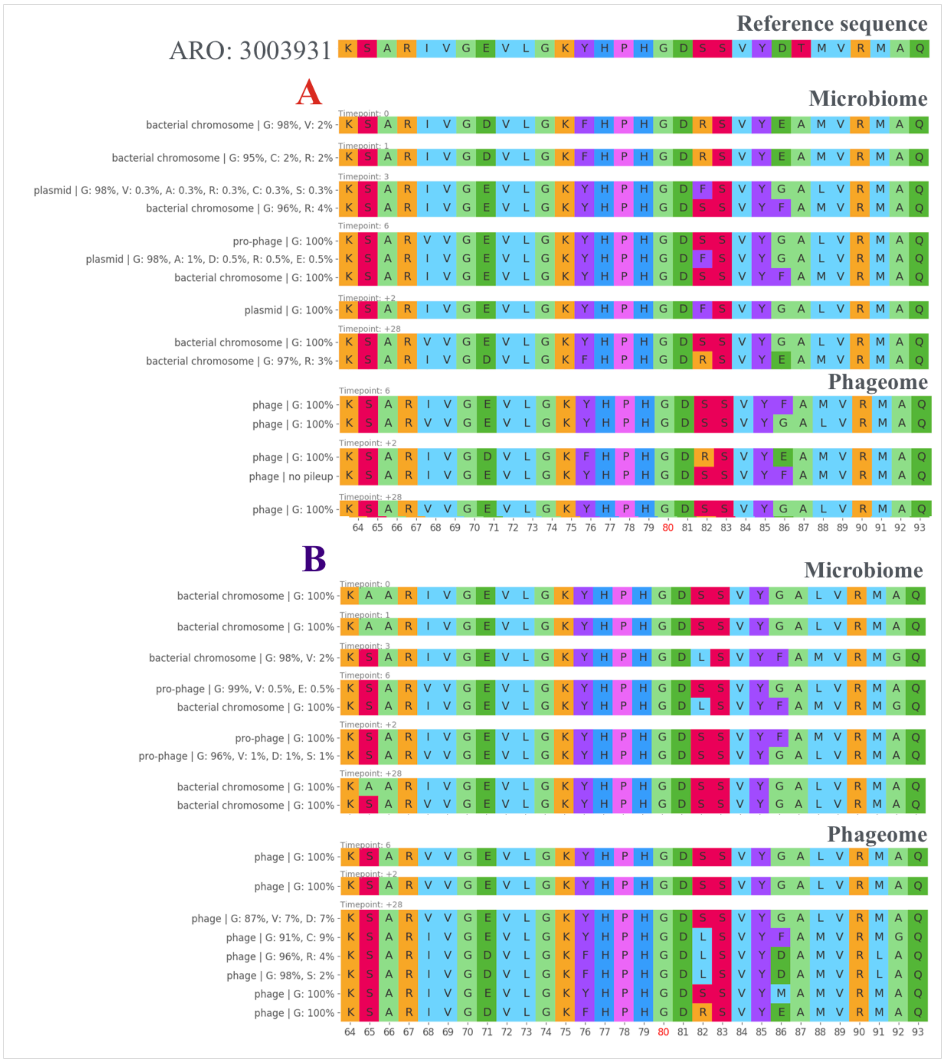


Figure S2: Alignment of the protein sequences of the genes annotated as gyrases conferring resistance to ciprofloxacin found on scaffolds for both sets and participants. The labels in the left side denote scaffolds assignment to the MGE group and portion of the reads carrying various amino acids.
